# Bivalent mRNA vaccine booster induces robust antibody immunity against Omicron subvariants BA.2, BA.2.12.1 and BA.5

**DOI:** 10.1101/2022.07.19.500616

**Authors:** Zhenhao Fang, Sidi Chen

**Affiliations:** Department of Genetics, Yale University School of Medicine, New Haven, CT, USA; System Biology Institute, Yale University, West Haven, CT, USA; Center for Cancer Systems Biology, Yale University, West Haven, CT, USA; Immunobiology Program, Yale University, New Haven, CT, USA; Molecular Cell Biology, Genetics, and Development Program, Yale University, New Haven, CT, USA; MD-PhD Program, Yale University, New Haven, CT, USA; Comprehensive Cancer Center, Yale University School of Medicine, New Haven, CT, USA; Stem Cell Center, Yale University School of Medicine, New Haven, CT, USA; Center for Biomedical Data Science, Yale University School of Medicine, New Haven, CT, USA

**Keywords:** COVID, variant adapted booster, bivalent mRNA vaccine, Omicron BA.5, BA.2, BA.2.12.1, Delta variant, lipid nanoparticle

## Abstract

As the immune protection conferred by first booster shot wanes over time and new Omicron subvariant emerges with stronger immune evasion, the need for variant-adapted COVID vaccine booster is increasingly imminent. However, the rapid replacement of dominant Omicron subvariants (from BA.1 to BA.2, then BA.2.12.1 and now BA.4/5) poses a great challenge to update COVID vaccine targeting the fast-evolving variants while maintaining potency against existing variants. It is a crucial question to ask which variant-based antigen(s) to use in the next generation COVID vaccine to elicit potent and broad response to past, present, and potential rising variants. Bivalent vaccine candidates have been under active clinical testing such as Modern mRNA-1273.214. In this study, we generate a Delta + BA.2 bivalent mRNA vaccine candidate and tested in animals. We compare the antibody response elicited by ancestral (wild type, WT), Delta, BA.2 spike based monovalent or Delta & BA.2 bivalent mRNA boosters against Omicron BA.2, BA.2.12.1 and BA.4/5 subvariants. In mice pre-immunized with two doses of WT lipid nanoparticle mRNA (LNP-mRNA), all three monovalent and one bivalent boosters elevated Omicron neutralizing antibody titers to various degree. The boosting effect of Delta and BA.2 specific monovalent or bivalent LNP-mRNAs is universally higher than that of WT LNP-mRNA, which modestly increased antibody titer in neutralization assays of Omicron BA.5, BA.2.12.1 and BA.2. The Delta & BA.2 bivalent LNP-mRNA showed better performance of titer boosting than either monovalent counterparts, which is especially evident in neutralization of Omicron BA.4 or BA.5. Interestingly compared to the neutralizing titers of BA.2 and BA.2.12.1 pseudovirus, BA.2 monovalent but not Delta & BA.2 bivalent booster suffered a significant loss of BA.4/5 neutralizing titer, indicative of broader activity of bivalent booster and strong neutralization evasion of Omicron BA.4 or BA.5 even in the BA.2 mRNA vaccinated individuals. These data provide evaluation of WT, Delta, BA.2 monovalent and bivalent boosters antibody potency against Omicron BA.2, BA.2.12.1 and BA.4/5 subvariants.

---

As the immune protection conferred by first booster shot wanes over time and new Omicron subvariant emerges with stronger immune evasion, the need for variant-adapted coronavirus disease (COVID) vaccine booster is increasingly imminent. On June 28, vaccine advisory committee of food and drug administration (FDA) voted in favor of updating COVID booster shot to add an Omicron component^1^. However, the rapid displacement of dominant Omicron subvariants (from BA.1 to BA.2, then BA.2.12.1 and now BA.4 and BA.5) poses a great challenge to update COVID vaccine targeting the fast-evolving variants while maintaining potency against circulating variants ^2^. Each former dominant Omicron strain, including BA.1, BA.2 and BA.2.12.1, drastically surges and subsides in a window of 3 months or even shorter^3^. Omicron BA.4 and BA.5 subvariants emerge in April in Southern Africa and become dominant around the world since June this year^3^. These Omicron sublineages quickly replace its predecessors in circumstances of existing herd immunity from vaccination or infection of past variants. Reinfection or breakthrough infection caused by new dominant variant is not uncommon due to its strong immune evasion^4, 5^, which complicates the redesign of new COVID boosters given the short time window of each Omicron wave and the lead time between design, validation and deployment of new boosters.

It is a crucial question to ask that which variant based antigen(s) to use in the next generation COVID boosters in order to elicit potent and broad response to past, present and emerging variants. At the time we initiated this study, the then-dominant subvariant BA.2 was gradually replaced by BA.2.12.1, BA.4 and BA.5. Compared to BA.2 spike, BA.2.12.1 contains two additional mutations (L452Q and S704L) while BA.4 and BA.5 spikes are identical and have 4 constant alterations (Del69-70, L452R, F486V, R493Q) plus one mutation (N658S) seen in earlier sequences (**Fig. 1a–1b**). The L452 substitutions in BA.2.12.1 and BA.4/5 are associated with neutralizing antibody escape^6^ and BA.4/5 combines the L452R mutation initially identified in Delta variant, highlighting one possible evolution trajectory of emerging variant by combining predecessors’ beneficial mutations.

**Figure 1.**
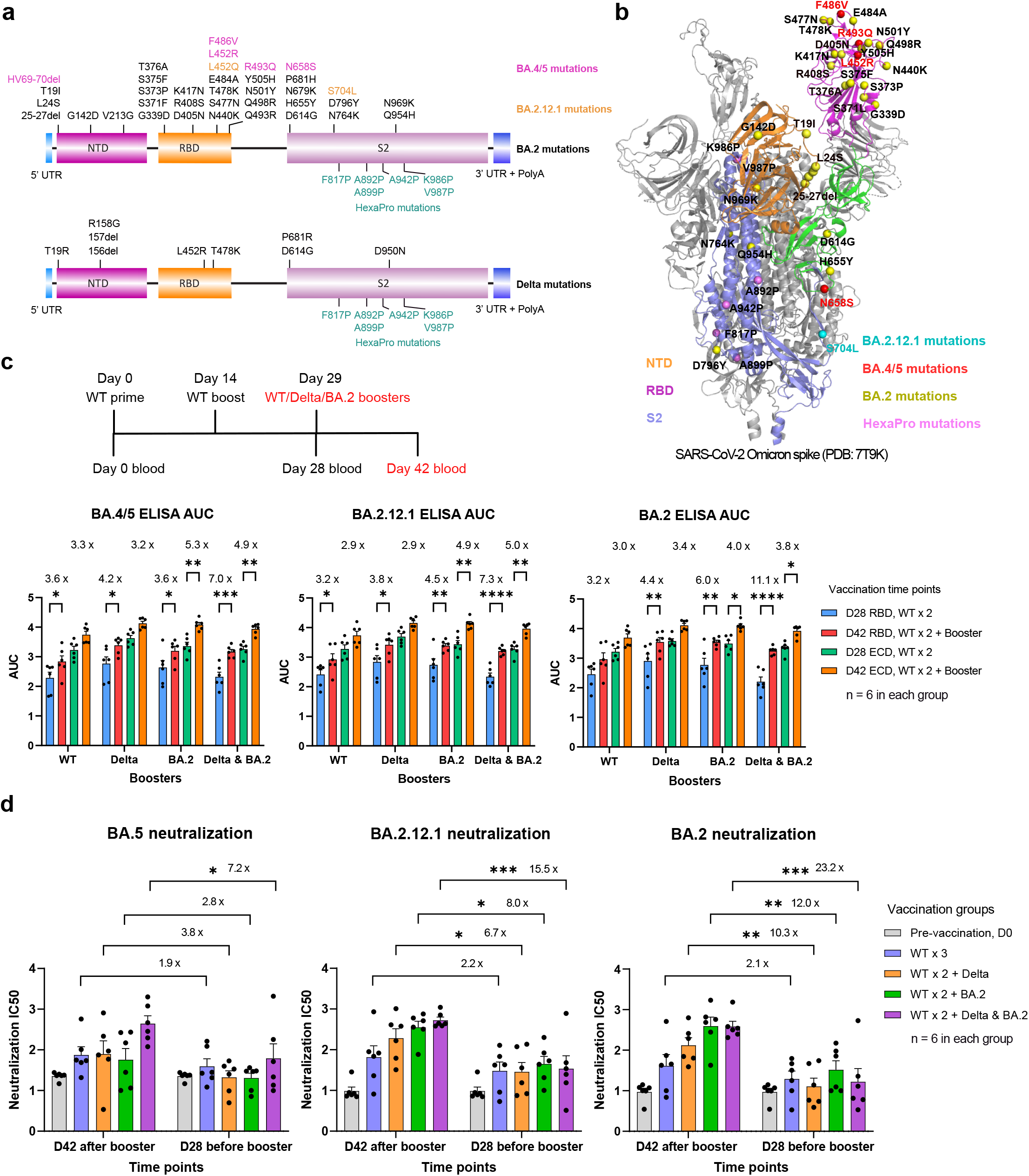
Potent antibody response to Omicron BA.2, BA.2.12.1 and BA.5 subvariants by Omicron BA.2 and Delta bivalent LNP-mRNA. **a**, Vaccine design of Omicron BA.2 and Delta variant specific LNP-mRNA based on BA.2 and Delta spike mutations. Unique spike mutations on BA.2.12.1 and BA.5 (not included in LNP-mRNA) are colored in orange and magenta. **b**, Distribution of BA.2 (Yellow), BA.2.12.1(Cyan) and BA.5 (Red) mutations in one protomer of Omicron spike trimer (PDB: 7T9K). **c**, Delta and BA.2 specific monovalent or bivalent LNP-mRNA boosters improved antibody response of WT-vaccinated mice to Omicron BA.2, BA.2.12.1 and BA.4/5 subvariants. Comparison of binding antibody titers against BA.2, BA.2.12.1 and BA.4/5 spike RBD and ECD before (D28) and after (D42) receiving 1.5 μg WT, Delta, BA.2 specific monovalent or bivalent (1.5 μg Delta + 1.5 μg BA.2) LNP-mRNA boosters. Antibody titers were quantified by area under curves (AUC) of ELISA response curves in Figure S1 and S2. Blood samples were collected in mice immunized with two doses of 1.5 μg WT LNP-mRNA followed by 1.5 μg WT, Delta, BA.2 specific monovalent or Delta & BA.2 bivalent boosters (n = 6 in each group). **d**, Neutralization of Omicron BA.2, BA.2.12.1 and BA.5 pseudovirus by plasma of mice before (D28) and after (D42) vaccinated with WT, Delta, BA.2 specific monovalent or Delta & BA.2 bivalent boosters. Six samples collected on day 0 were included and compared to both D28 and D42 datasets. Titer ratios before and after receiving boosters (D42/D28 ratios) were shown in c-d. Individual dot in dot-bar plots represent value from each mouse and are shown as mean ± s.e.m.. To assess statistical significance, two-way ANOVA with Tukey’s or Šídák’s multiple comparisons test was used. Statistical significance labels: * p < 0.05; ** p < 0.01; *** p < 0.001; **** p < 0.0001. Non significant comparisons are not shown.

Bivalent vaccine candidates have gained recent tractions due to the concept of direct targeting of two variants, which may also induce broader immunity against other variants. Bivalent vaccine candidates have been under active clinical testing such as Modern’s mRNA-1273.214, which is a equal mixture of two spike-encoding mRNAs with 25 μg targeting ancestral SARS-CoV-2 and 25 μg targeting the original Omicron Variant (B.1.1.529) (Moderna news releases June 08 2022, June 22 2022, and FDA committee meeting June 28 2022), demonstrating the importance and the clinically relevance of the concept of bivalent vaccination using two mRNAs. In light of this merge of variants’ mutations (**Fig. 1a–1b**), we want to ask if mRNA vaccine candidates based on antigens of circulating variant (BA.2) and/or former dominant variant (Delta) can mediate broad antibody response to emerging variants such as BA.2.12.1, BA.4 or BA.5. It is worth to explore in this direction for a few reasons. The lead time of combining boosters adapted to dominant and former dominant variants will be shorter than predicting and developing boosters targeting new variants. In addition, because of the rapid displacement of circulating variants, the mismatch between the strain used for updated boosters and emerging strain may always exists. How to elicit broad response to emerging variants using existing variant antigens is an inevitable question to answer when redesigning updated COVID boosters.

To answer this question, we compared the antibody response elicited by ancestral (wild type, WT), Delta, BA.2 spike based monovalent or Delta & BA.2 bivalent mRNA boosters against Omicron BA.2, BA.2.12.1 and BA.4/5 subvariants. In mice pre-immunized with two doses of WT lipid nanoparticle mRNA (LNP-mRNA), all three monovalent and one bivalent boosters elevated Omicron binding and neutralizing antibody titers to various degree in ELISA and pseudovirus neutralization assay (**Fig. 1c–1d and Figs. S1–S3**), exemplifying the benefit of receiving WT or variant-adapted booster shots against circulating and emerging variants. Booster-associated titer ratios quantify booster’s effect on antibody titers and were shown in each bar graph as post-booster titer on day 42 over pre-booster titer on day 28. Its dynamic range was greater in neutralization assay (ratio ranges from 3-23) than in ELISA (ratio ranges from 2-11).

Before administered with different boosters, 24 mice in four groups received same treatment and showed little or no significant difference in antibody titers measured on day 0 and day 28 (**Figs. S4–S6 and S7a**). A moderate increase in Omicron neutralizing antibody titers was observed from immunization of two doses of WT LNP-mRNA (**Fig. S7b**). This titer increase by WT LNP-mRNA was lowest in neutralization assay of BA.4/5 (~40% increase) as compared to BA.2.12.1 and BA.2. On day 42 two weeks post booster, the binding and neutralizing titers of WT booster group were frequently found lower than those of variant booster groups (**Fig. S4 and S7a**), consistent with the fact that BA.4/5 have stronger evasion of existing antibody therapeutics or vaccine induced immunity. Interestingly, compared to the neutralizing titers of BA.2 and BA.2.12.1, BA.2 monovalent but not Delta & BA.2 bivalent booster suffered a significant loss of BA.4/5 neutralizing titer (**Fig. S7c**), indicative of broader activity of bivalent booster and strong neutralization escape of Omicron BA.4 or BA.5 even in the BA.2 mRNA vaccinated individuals. The RBD and ECD binding antibody titers were well correlated and showed distinct linear regression models between day 28 and day 42 as well as in WT, Delta (right panel in **Fig. S5**) and Omicron antigen datasets (left panel). The upper right shift of day 42 linear segment suggested a titer increase by boosters while the lower left shift in Omicron antigen dataset was associated with antibody evasion of Omicron antigens.

The boosting effect of Delta and BA.2 specific monovalent or bivalent LNP-mRNAs is universally higher than that of WT LNP-mRNA, which only modestly increased antibody titer (~1 fold, fold change = ratio – 1) in neutralization assays of Omicron BA.5, BA.2.12.1 and BA.2 (**Fig. 1d**). The Delta & BA.2 bivalent booster showed superior performance of enhancing binding and neutralizing titers than either monovalent counterparts, which is especially apparent in neutralization of Omicron BA.4 or BA.5. The bivalent booster associated titer ratios were 23, 16 and 7 fold for neutralization of BA.2, BA.2.12.1 and BA.4/5, respectively while Delta/BA.2 monovalent booster ratios were 10/12, 7/8, 4/3 respectively. The linear regression models of neutralizing and binding titers showed a trend of correlation but the goodness of fit was low due to deviations intrinsic in the two assays as well as heterogeneity stemmed from distinct boosters and Omicron subvariants tested (**Fig. S8**).

To sum up, our data delivered a few clear messages regarding the potency of boosters against Omicron subvariants: 1) either WT or variant, monovalent or bivalent boosters can improve antibody response to Omicron BA.2, BA.2.12.1 and BA.4/5, demonstrating the benefit and necessity of receiving booster shots; 2) the variant boosters with closer antigenic distance to circulating variant perform universally better than WT booster; 3) compared to monovalent booster, bivalent booster combining two genetically distant variants, Delta & BA.2 showed broader and numerically stronger antibody response to Omicron BA.2, BA.2.12.1 and BA.4/5 subvariants. Taken together, our study presents a direct evaluation of Delta and BA.2 variant-adapted monovalent and bivalent mRNA boosters and compares their antibody response to Omicron subvariants with WT booster in the context of mouse model pre-immunized with two-dose WT LNP-mRNA vaccination. These data provide pre-clinical evidence and rationale for developing bivalent or multi-valent variant targeted COVID boosters.

## Acknowledgement

This work is supported by DoD PRMRP IIAR (W81XWH-21-1-0019) and discretionary funds to S.C. We thank members from our labs for support. We thank support from Institutes of Systems Biology and Cancer Biology; Yale core facilities, Department of Genetics; Dean’s Office of Yale School of Medicine and the Office of Vice Provost for Research.

## Supplementary figure legends

**Figure S1.**
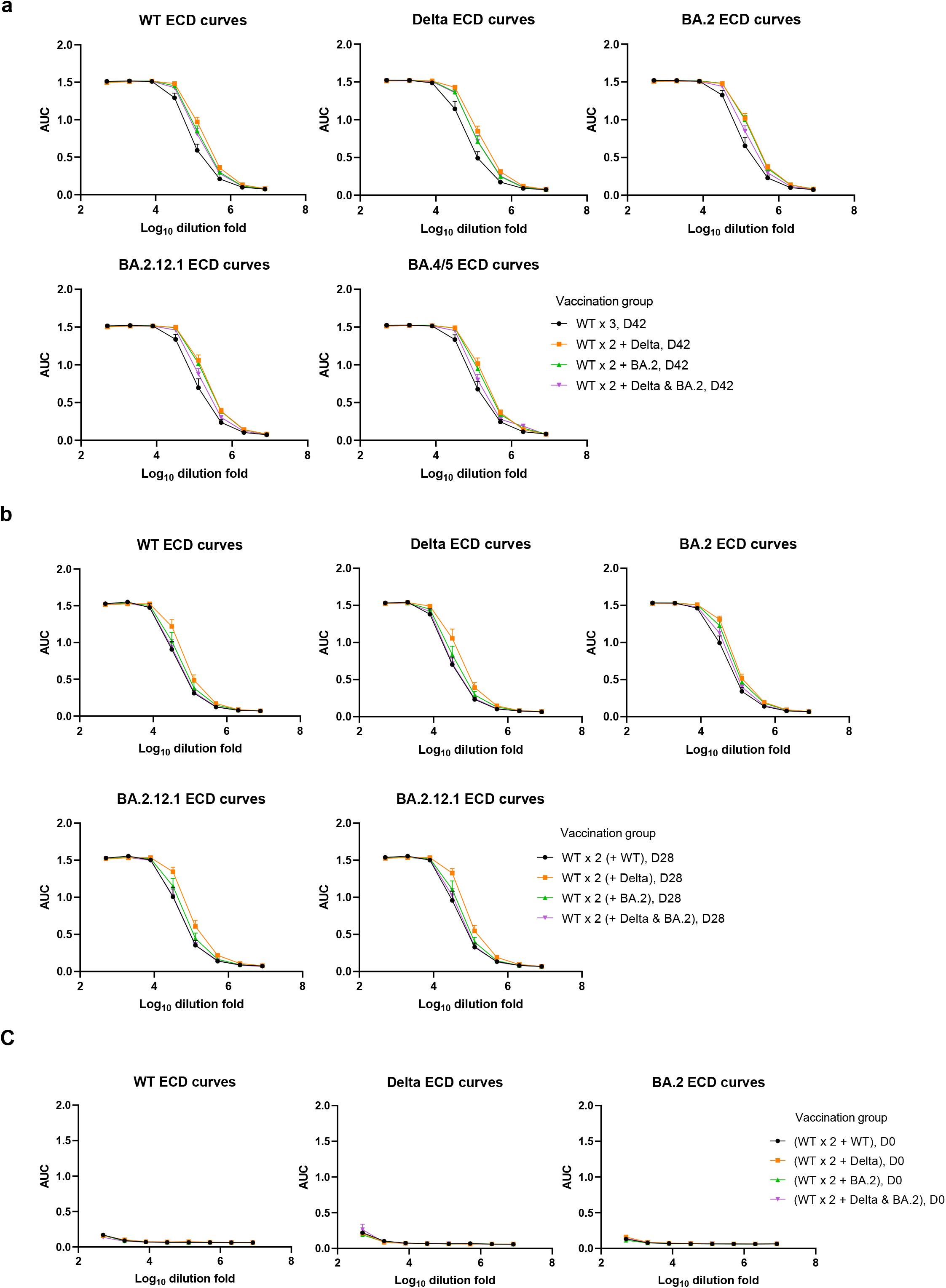
Plasma dilution-dependent ELISA response curves against WT, Delta, BA.2, BA.2.12.1 and BA4/5 spike ECDs. Plasma samples were collected at day 42 (a), day 28 (b) and day 0 (c) from mice immunized with WT Delta, BA.2 specific monovalent or bivalent LNP-mRNA boosters

**Figure S2.**
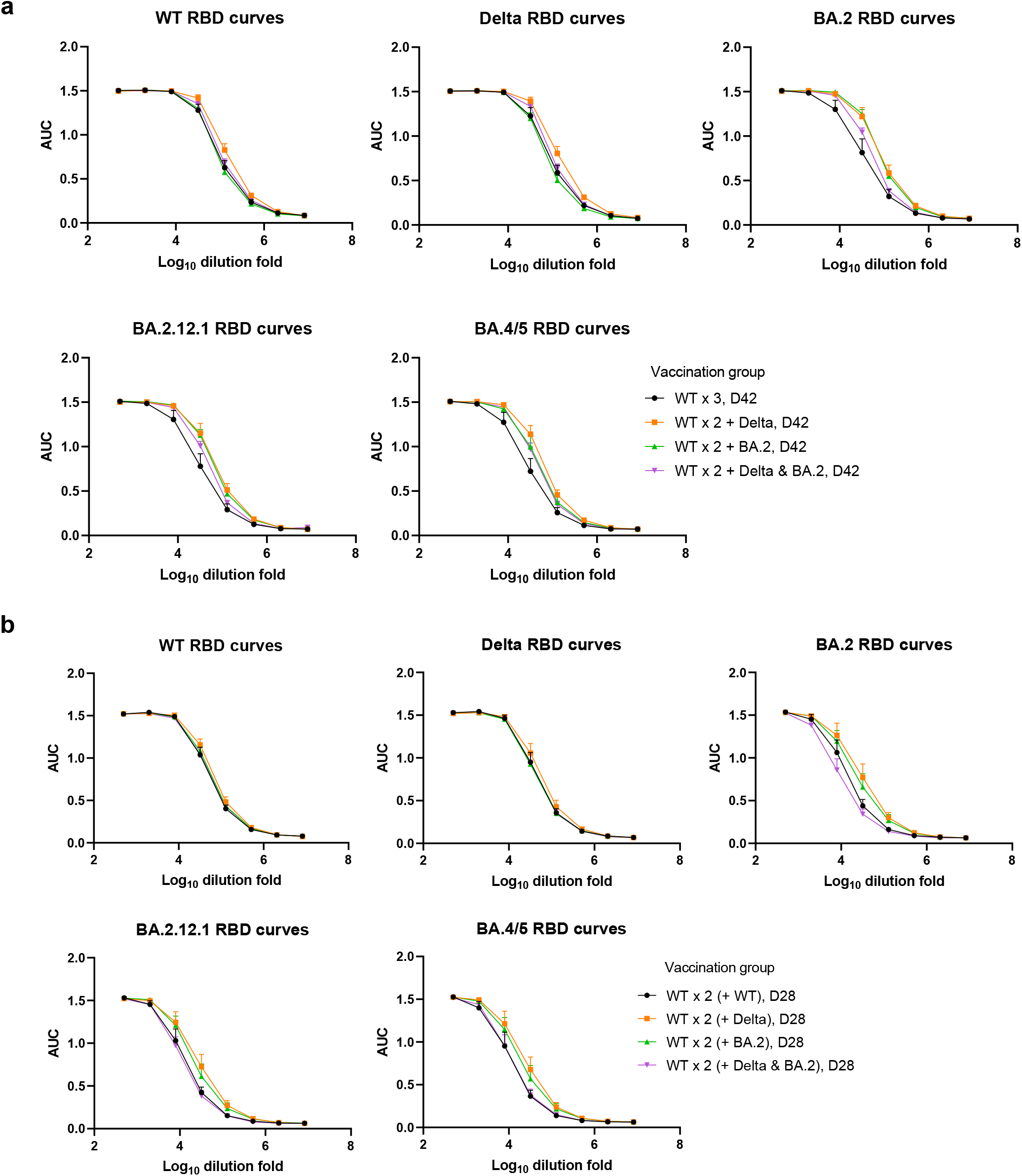
Plasma dilution-dependent ELISA response curves against WT, Delta, BA.2, BA.2.12.1 and BA4/5 spike RBDs. Plasma samples were collected at day 42 (a) and day 28 (b) from mice immunized with WT Delta, BA.2 specific monovalent or bivalent LNP-mRNA boosters.

**Figure S3.**
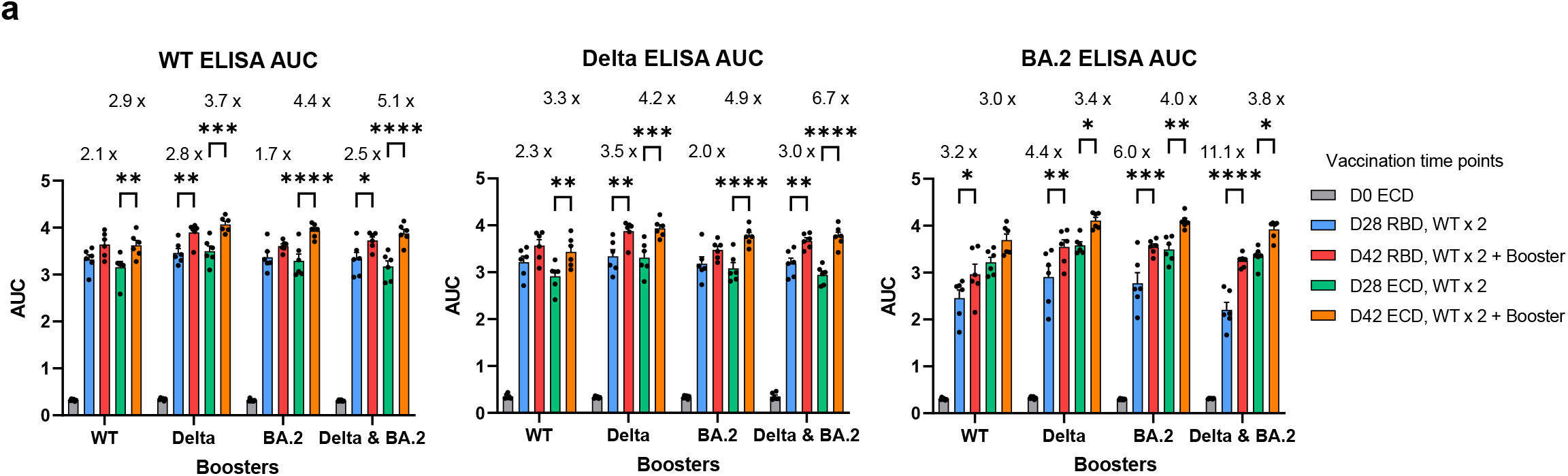
Comparison of binding antibody titers against WT (left), Delta (Mid) and BA.2 (Right) spike RBD and ECD before (D0 and D28) and after (D42) receiving 1.5 μg WT, Delta, BA.2 specific monovalent or bivalent (1.5 μg Delta + 1.5 μg BA.2) LNP-mRNA boosters (n = 6). Antibody titers were quantified by area under curves (AUC) of ELISA response curves in Figure S1 and S2. The comparison with day 0 samples and insignificant comparison were not shown.

**Figure S4.**
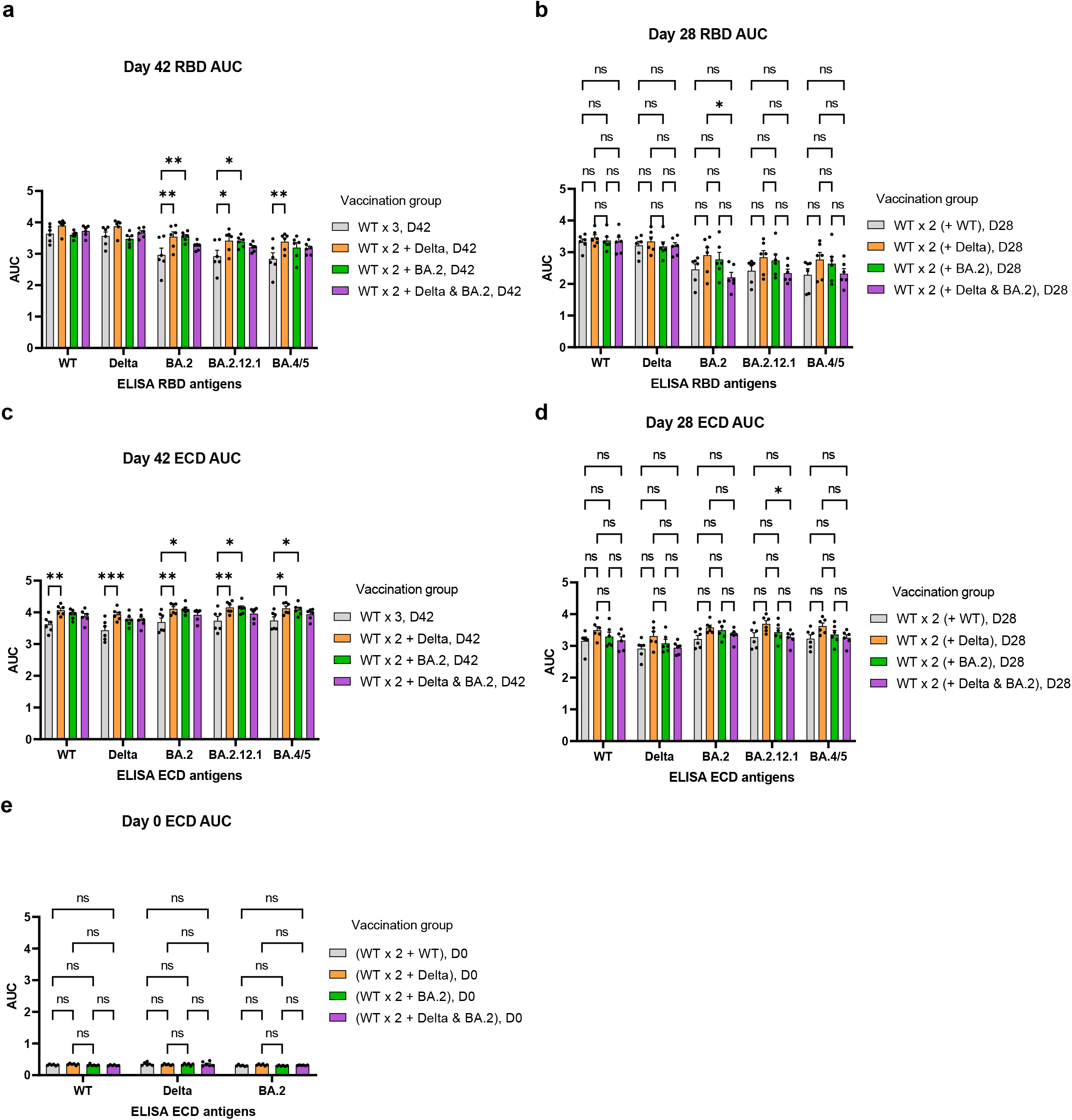
Comparison of ELISA antibody titers of plasma samples collected on day 0, day 28 and day 42. **a-b**, ELISA antibody titers against WT, Delta, BA.2, BA.2.12.1 and BA.4/5 spike RBDs before (D28, b) and after (D42, a) receiving 1.5 μg WT, Delta, BA.2 specific monovalent or bivalent (1.5 μg Delta + 1.5 μg BA.2) LNP-mRNA boosters. **c-e**, ELISA antibody titers against WT, Delta, BA.2, BA.2.12.1 and BA.4/5 spike ECDs by plasma samples collected on (D42, c; D28, d; D0, e). Antibody titers were quantified by area under curves (AUC) of ELISA response curves in Figure S1 and S2.

**Figure S5.**
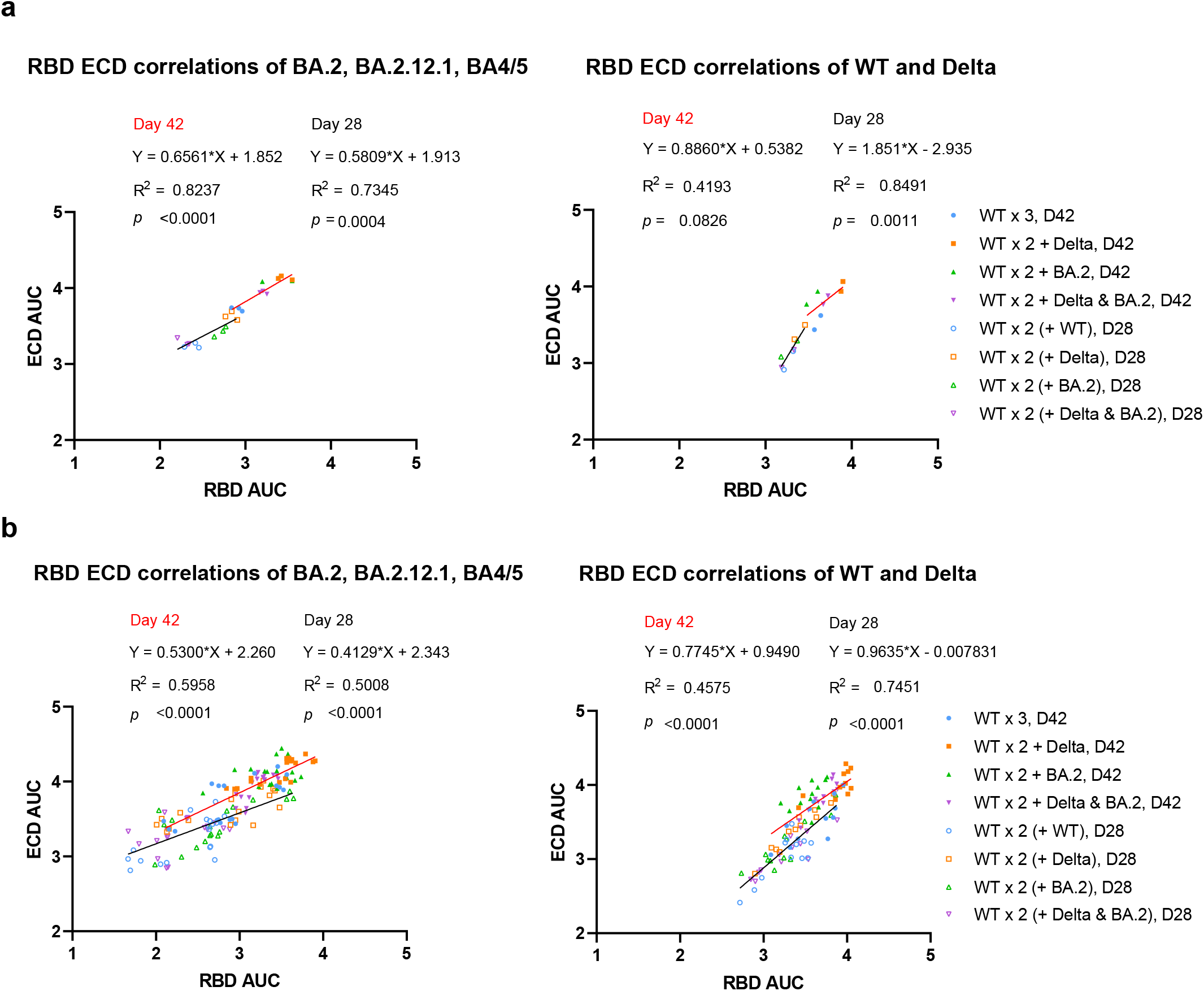
Correlation of antibody titers against RBD and ECD of five spike antigens in ELISA. Antibody titers against ECD of Omicron BA.2, BA.2.12.1, BA.4/5 subvariants (left) or WT, Delta (right) were shown on y axis as log_10_ AUC and plotted against corresponding RBD binding antibody titers on x axis (log_10_ AUC). Titers were either shown as mean of matched vaccination group (a) or derived from individual animal (b).

**Figure S6.**
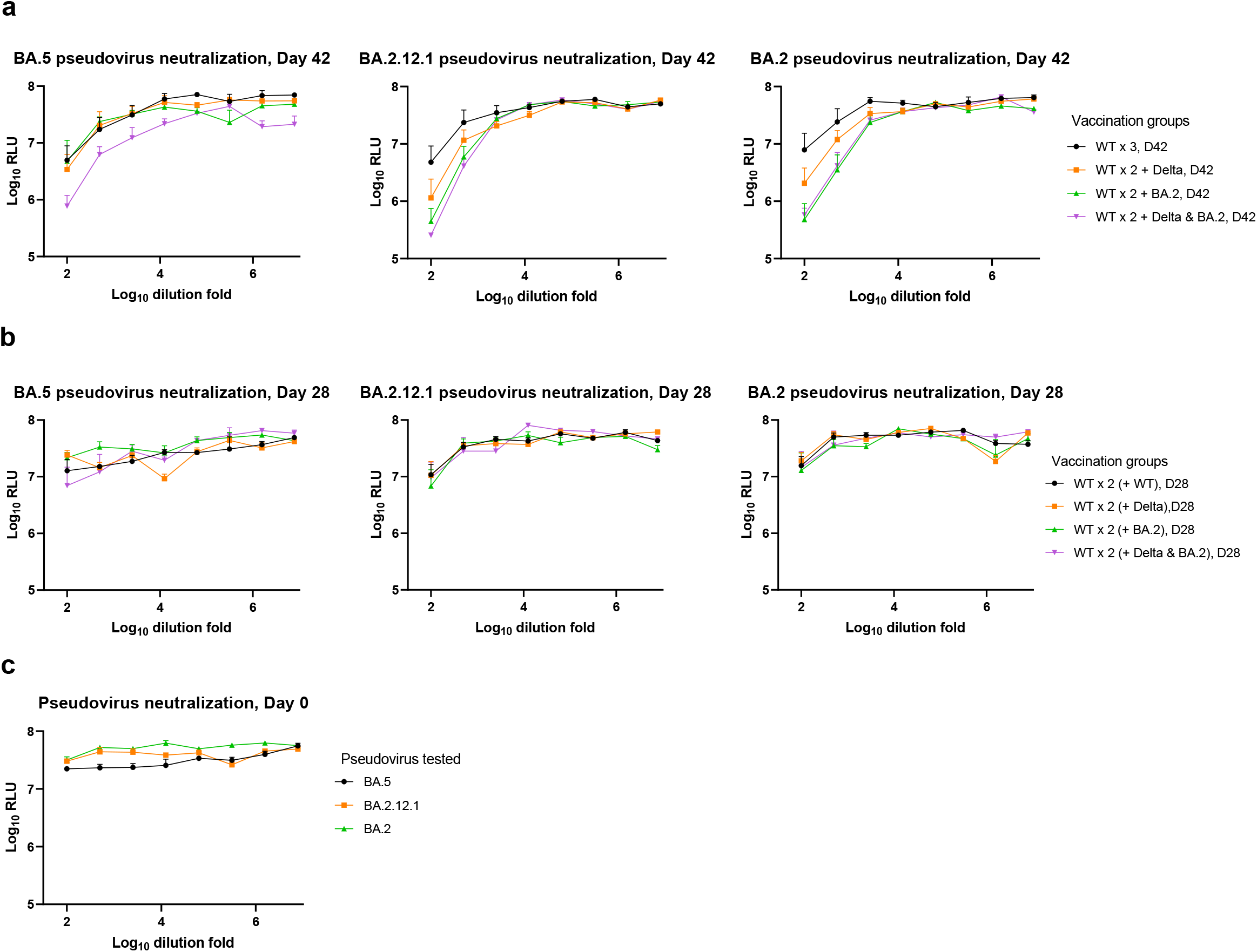
Neutralization titration curves of serially diluted plasma collected at indicated time points from mice vaccinated with WT, Delta, BA.2 monovalent or bivalent LNP-mRNA boosters. **a,** Neutralization curves of BA.5, BA.2.12.1 and BA.2 pseudovirus by samples collected on day 42 from mice immunized with 1.5 μg WT, Delta, BA.2 monovalent or bivalent LNP-mRNA boosters. **b**, Neutralization curves of BA.5, BA.2.12.1 and BA.2 pseudovirus by samples collected on day 28 from mice immunized with two doses of 1.5 μg WT LNP-mRNA. **c**, Neutralization curves of BA.5, BA.2.12.1 and BA.2 pseudovirus by samples collected on day 0 from vaccination naïve mice. The log_10_ relative light unit (RLU) measured by NanoLuc luciferase assay were shown as mean ± s.e.m. and plotted against serial log10-transformed sample dilution points.

**Figure S7.**
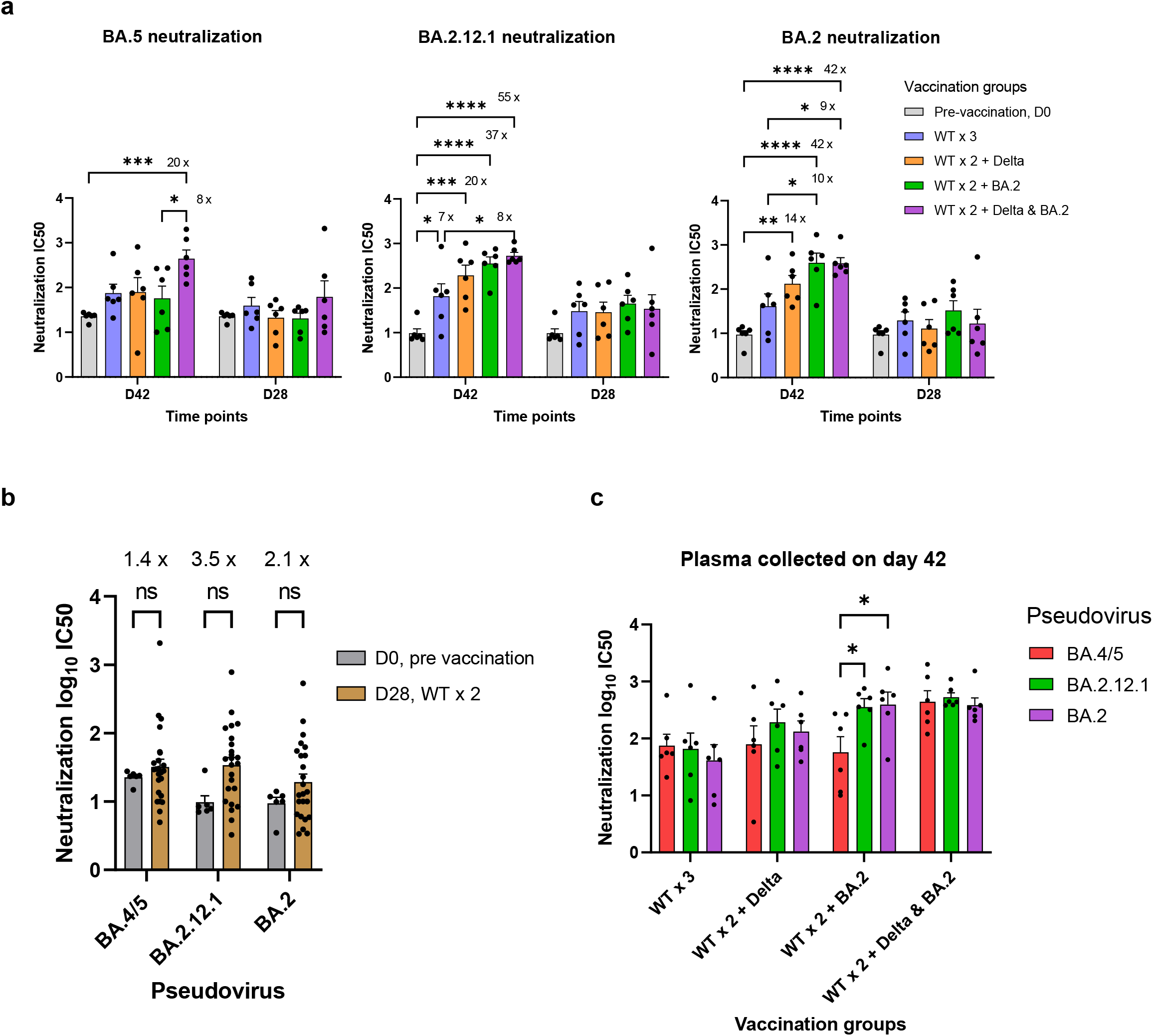
Statistical comparison of neutralizing titers of plasma samples from different vaccination groups at same time point (a) or against different Omicron subvariant pseudoviruses at matched time points (b). **a**, Omicron BA.2 (right), BA.2.12.1 (mid) and BA.5 (left) pseudovirus neutralization by plasma of mice before (D28) and after (D42) vaccinated with WT, Delta, BA.2 specific monovalent or Delta & BA.2 bivalent boosters. Six samples collected on day 0 were included and compared to both D28 and D42 datasets. **b**, BA.4/5, BA.2.12.1 and BA.2 neutralizing antibody titers from samples collected on day 0 and day 28 (WT x 2) were compared. **c**, BA.4/5, BA.2.12.1 and BA.2 neutralizing antibody titers were compared within same vaccination groups at matched time points including day 28 (pre booster) and day 42 (post booster).

**Figure S8.**
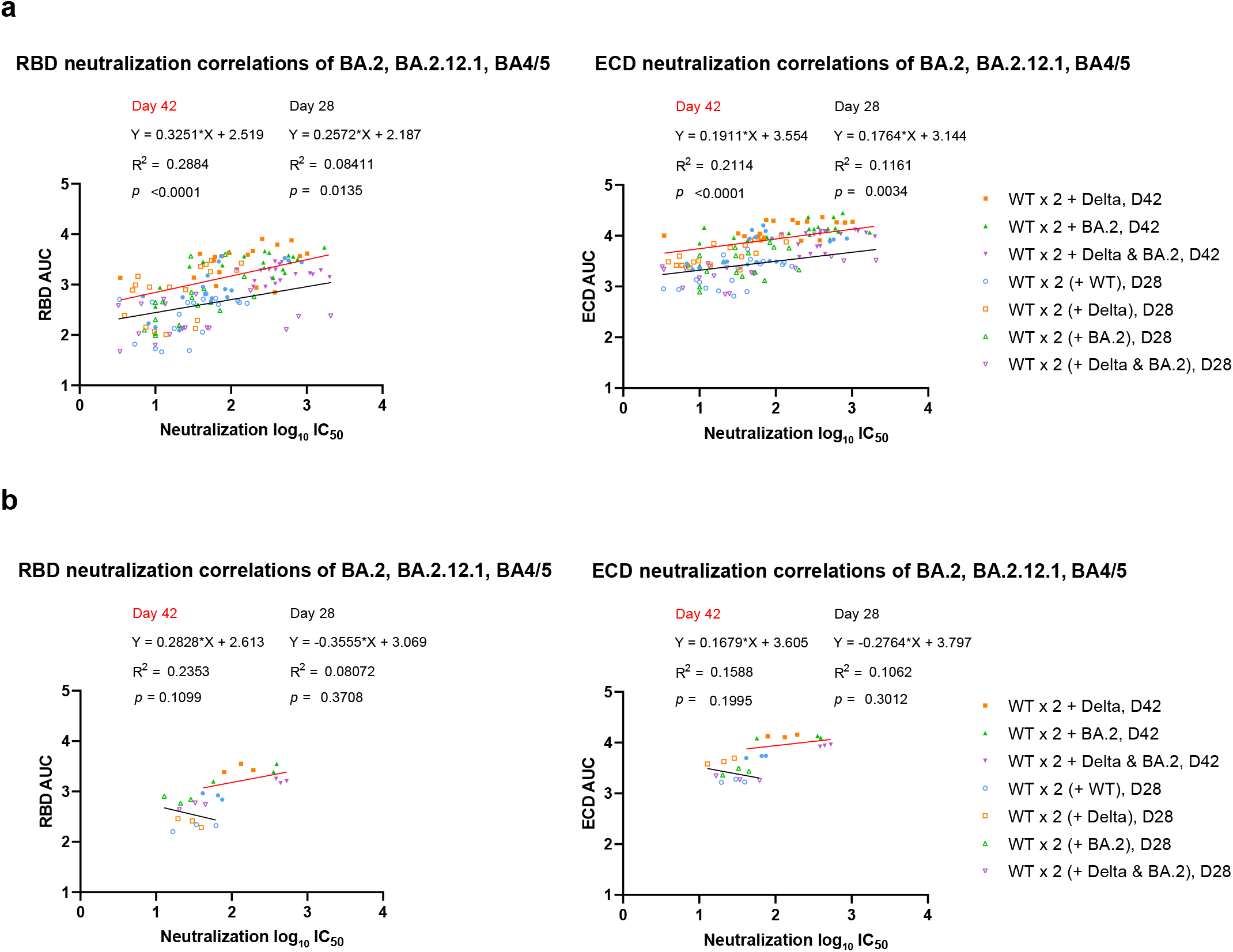
Correlation of antibody titers measured by pseudovirus neutralization and ELISA. Antibody titers determined by pseudovirus neutralization assay were shown on x axis as log_10_ IC50 and plotted against ELISA binding antibody titers (log10 AUC) measured by RBD (left) or ECD (right) spike antigens on y axis. Titer values were either derived from mean of matched vaccination group (b) or individual animals (a).

## Methods

### Institutional approval

All animal work was performed under the guidelines of Yale University Institutional Animal Care and Use Committee (IACUC) with approved protocols (Chen 2020-20358; Chen 2021-20068; Wilen 2021-20198). All recombinant DNA (rDNA) and biosafety work were performed under the guidelines of Yale Environment, Health and Safety (EHS) Committee with approved protocols (Chen 18–45, 20–18, and 20–26).

### Molecular cloning and mRNA preparation

The WT and Delta spike plasmids were cloned in our previous study^7, 8^. BA.2 spike plasmid was cloned based on the isolate sequencing data in GISAID EpiCoV (EPI_ISL_6795834.2)^9^. WT, Delta and BA.2 spike plasmids were linearized by restriction enzymes and transcribed to mRNA by in vitro T7 RNA polymerase (NEB, Cat # E2060S) as previously described^10^.

### Cell culture

hACE2-293FT and 293T cells were cultured in Dulbecco’s minimal essential medium (DMEM, Fisher) supplemented with 10% fetal bovine serum (Hyclone) and penicillin (100 U/ml)-streptomycin (100 ug/ml). Cells were split ever other day at a 1:4 ratio when confluency is over 90%.

### Lipid nanoparticle mRNA preparation

In brief, lipids mixture was solubilized in ethanol and mixed with spike mRNA in pH 5.2 sodium acetate buffer. The mRNA encapsulated by LNP (LNP-mRNA) was then buffer exchanged to PBS using 100kDa Amicon filter (Macrosep Centrifugal Devices 100K, 89131-992). The size distribution of LNP-mRNA was evaluated by dynamic light scatter (DynaPro NanoStar, Wyatt, WDPN-06). The Quant-iT™ RiboGreen™ (Thermo Fisher) RNA Assay was applied to determine encapsulation rate and mRNA amount.

### Animal vaccination

Animal immunization was performed on 16-18 weeks female C57BL/6Ncr mice purchased from Charles River. Mice were vaccinated with two doses of 1.5 μg WT LNP-mRNA on day 0 and day 14 followed by 1.5 μg WT, Delta, Omicron BA.2 monovalent booster or Delta & BA.2 bivalent booster on day 29. The plasma samples were isolated from blood which was collected before vaccination on day 0, two weeks after WT boost on day 28 and two weeks after monovalent or bivalent boosters on day 42.

### ELISA and Neutralization assay

The binding and neutralizing antibody titers were determined by ELISA and pseudovirus neutralization assay as previously described^10^. NanoGlo luciferase assay system (Promega N1120) was applied to determine the pseudovirus infection level in hACE2-293FT cells. The ELISA antigens including RBDs of WT (Sino 40592-V08B), Delta(Sino 40592-V08H90), Omicron BA.2(Acro SPD-C522g-100ug), BA.2.12.1(Acro SPD-C522q-100ug) and BA.4/5(Acro SPD-C522r-100ug) were purchased from Sino Biological and AcroBiosystems. The ELISA ECD antigens including WT (Sino 40589-V08B1), Delta (Sino 40589-V08B16), Omicron BA.2 (Acro SPN-C5223-50ug), BA.2.12.1 (Acro SPN-C522d-50ug) and BA.4/5 (SPN-C5229-50ug) were purchased from Sino Biological and AcroBiosystems. The pseudovirus plasmids of spike without HexaPro mutations were generated based on the WT plasmid which was a gift from Dr. Bieniasz’s lab.

### Data availability

All source data and statistical analysis are provided in this article and its supplementary excel file.

### Code availability

No custom code was used in this study.

